# Muscle Strength and Size Relationships with Unilateral Progressive Resistance Training

**DOI:** 10.1101/2025.01.13.632853

**Authors:** Donald D. Deiwert, Sisi Ma, Christopher Carey, Davin Greenwell, Heather Gordish-Dressman, Paul D. Thompson, Thomas Price, Theodore J. Angelopoulos, Priscilla M. Clarkson, Paul M. Gordon, Niall M. Moyna, Linda S. Pescatello, Paul S. Visich, Robert F. Zoeller, Eric P. Hoffman, Monica J. Hubal

**Affiliations:** Department of Kinesiology, Indiana University Indianapolis, Indianapolis IN; Institute for Health Informatics, University of Minnesota, Minneapolis MN; Department of Genomics and Precision Medicine, George Washington University, Washington DC; Department of Cardiology, Hartford Hospital Connecticut, Hartford CT; College of Health Sciences, University of Bridgeport, Bridgeport CT; Department of Rehabilitation & Movement Science, University of Vermont, Burlington VT; Department of Health, Human Performance, & Recreation, Baylor University, Waco TX; Department of Clinical Exercise Physiology, Dublin City University, Whitehall, Dublin, Ireland; Department of Kinesiology, University of Connecticut, Storrs CT; Department of Exercise and Sport Performance, University of New England, Biddeford ME; Department of Exercise Science and Health Promotion, Florida Atlantic University, Bocca Raton FL; Department of Pharmaceutical Sciences, Binghamton University, Binghamton NY

**Keywords:** causal modeling, heterogeneity, resistance training, sex differences

## Abstract

**Purpose:** This study defines correlative and causal relationships between muscle strength and size before and after unilateral resistance training (RT) in a large cohort of healthy adults, focusing on sex differences within these relationships.

**Methods:** Results from 1233 participants (504 males and 729 females) in a retrospective analysis were included. Maximal voluntary isometric contraction strength (MVC), one-repetition maximum strength (1RM), biceps cross-sectional area (CSA) and elbow flexor volume (VOL) measures of the non-dominant and dominant arm were evaluated from baseline and after 12-wk RT twice per week. Correlations of MVC and VOL and 1RM and VOL were calculated in the whole cohort and within each sex independently. Causal analysis modeling was used to infer mechanistic relationships among variables.

**Results:** Absolute muscle strength and size related to one another both at baseline and following training, however correlation strength in each sex were weak. After RT, MVC relative change and VOL relative change correlations were correlated for the whole cohort (r=0.16; p<0.001) and females (r=0.18; p<0.001), but not in males (r=0.11; p=0.07). No significant correlations for relative change in 1RM and VOL were observed for the whole cohort or within sex. Causal discovery determined that change in VOL caused significant change in 1RM (but not MVC) and age was identified as a potential cause.

**Conclusions:** Sex differences occur in muscle size and strength relationship adaptations following resistance training, most notably the absence of significant relationships between relative size and strength changes in men. Simpson’s paradox bias, where assessing the combined data of males and females (also affecting overall sample size) affects identifies patterns differently than assessing relationships within each sex, may partially explain our findings.

## Introduction

Convention has long held the precept that skeletal muscle size is directly linked to its function. This perception is rooted in a phylogenetic tendency towards appraising larger individuals as stronger (1). It may also be a result of the common and overlapping training modalities by which increases in muscle strength and muscle size are elicited (2, 3). Excluding the elastic properties inherent to skeletal muscle and connective tissue, muscle force production is generally attributed to forces from myofibrillar actin-myosin cross bridging (4), which is a complex product of neural input patterns, existing structural architecture and metabolic capacity. As muscle produces force, it stands to reason that a larger muscle produces greater force. Linking muscle size and strength is not abjectly wrong. Rather, evidence is mounting suggesting that the factors underlying the relationship between muscle size and strength and their interdependence are more nuanced than some think, especially with interventions like resistance training.

Resistance training can induce both muscle hypertrophy and changes in muscle contractile properties (5-7). However, the degree to which each of these changes are elicited is tied to the specific training parameters employed (2, 8). Variations in task demands (load, timing, positioning, repetition, etc.) stress these underlying components of muscle strength in unique ways, and the specificity of training principle states that the unique demands of a given training program will elicit changes specific to the training stimuli. Muscle size adaptations also occur with overload training and are dependent on dynamics of underlying processes like alterations in the balance of muscle protein synthesis and breakdown (protein turnover) and stimulation and incorporation of muscle satellite cells to increase fiber size (9).

Debates have argued to what extent strength and size increases in response to resistance training are related (10, 11). However, the studies cited by Folland and Balshaw (10) that utilized high-resolution magnetic resonance imaging (MRI) and supported the relationship all suffered from relatively small sample sizes (< 40 participants) (12-14). These small sample sizes could undermine the findings due to being underpowered, especially considering the high inter-subject variation in response to resistance training as demonstrated by (15).

In the largest resistance training study published to date, we (15) previously demonstrated sex-dependent wide response ranges in strength and size changes following a standardized 12-week progressive unilateral resistance training program involving >500 untrained participants. Muscle size changes (% change) ranged from -2.5 to 55.5% in and -2.3 to 59.3% in females. Isometric strength changes (MVC) ranged from -24.3 to 148.5% in males and -31.5 to 93.4% in females. Dynamic strength (1RM) changes ranged from 0 to 150% in males and 0 to 250% in females. Despite there being overlap in these ranges, males demonstrated significantly greater muscle cross-sectional area (CSA % change) than females (20.4% vs. 17.9%). However, females demonstrated significantly greater relative change for isometric strength (22.0% vs. 15.8%) and 1RM strength (64.1% vs. 39.8%). While this study (15) provided insight on the variability and potential sex differences that occur with resistance training, the study did not answer whether males and females differ in the nature of the relationship <u>between</u> strength and size variables.

The present study’s purpose was to further elucidate muscle strength and muscle size changes by assessing relationships between strength and size in a very large cohort of untrained young healthy adults at baseline and following 12 weeks of unilateral progressive resistance training of the non-dominant elbow flexors/extensors. The unilateral intervention design allowed for the dominant arm to serve as a non-exercised control. We hypothesized that size and strength parameters would be significantly associated at baseline and following training, but that the relationships in change in size or strength would be more highly varied, demonstrating the complexity of adaptation. We also hypothesized that sex differences would be present in the strength/size relationships.

## Methods

### Study Overview

The current study uses de-identified data derived from a previously completed parent study (15, 16). Figure 1 displays a study overview. Main outcome measures (strength and size) were assessed at baseline and after 12-weeks of unilateral resistance training. The elbow flexor strength measures included isometric maximum voluntary contraction (MVC) and 1-repetition maximum (1RM). Whole-muscle upper arm size measures from MRI included two-dimensional CSA and three-dimensional whole-muscle volume (VOL).

**Figure 1.**
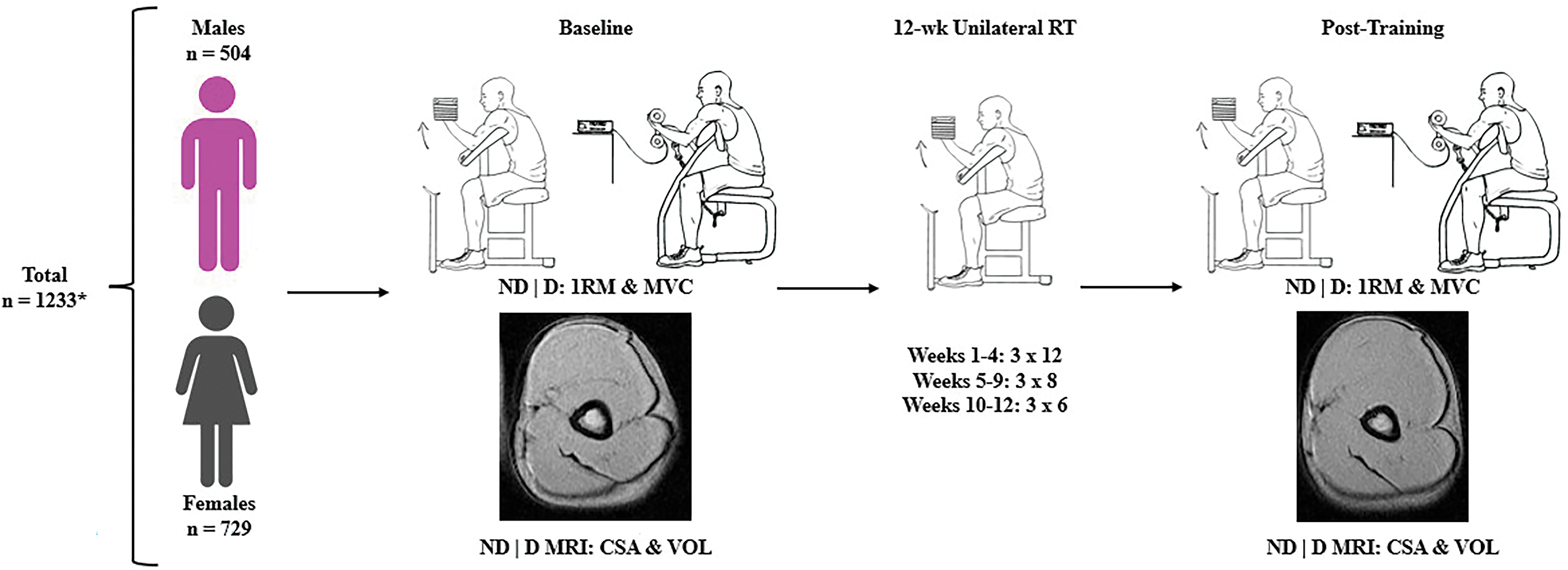
Overview of study. * Not all participants who were recruited had complete data. ND = non-dominant arm. D = dominant arm. MVC = maximum voluntary contraction. 1RM = 1-repetition maximum. MRI = magnetic resonance imaging. CSA = cross-sectional area. VOL = volume. RT = resistance training.

### Participants

Healthy, untrained males (n = 504) and females (n = 729) between 18-40 years were recruited for this study. This represents an addition of 648 subjects to the cohort previously described (15). Participant data were collected from 2002 to 2005 across a consortium of eight study centers (University of Massachusetts, Hartford Hospital, University of Connecticut, Florida Atlantic University, West Virginia University, University of Central Michigan, Dublin City University and, University of Central Florida) and sent to Children’s National Medical Center (CNMC) as a central processing site. MRI data were processed at Yale University and CNMC. Each site maintained approvals from their Institutional Review Board throughout the study and participants were informed of the study and read and signed an informed consent document. All data for the current analytics were de-identified.

In the complete cohort, ∼76% identified as Caucasian whereas the other ∼24% was made up of those identifying as Asian (∼36%), Hispanic (∼24%), African American (∼21%), “Other” (∼19%), and American Indian (∼1%). Exclusion criteria included consuming on average >2 alcoholic drinks daily and recent history of resistance training or resistance-training-like employment. Participants were also excluded from the study if they consumed any products that could affect skeletal muscle hypertrophy, such as androgenic precursors and protein supplements. Participants were instructed to maintain current dietary patterns and body weight.

### Muscle Strength Measures

MVC testing was performed using a modified preacher bench connected to a strain gauge (model 32628CTL, Lafayette Instrument Company, Lafayette, IN) as previously described (15). Baseline MVC measures at 90 deg elbow angle were gathered over three days (allowing at least 48h between days for recovery). On each day, three trials were completed with 60 seconds rest between attempts. Subjects were familiarized with testing on baseline day one, and the average measures from the next two days were considered the baseline MVC value. Baseline 1RM testing was only collected on baseline day three due to the protocol’s acute effect on muscle function, and it was performed using dumbbells while subjects were seated on a preacher curl bench. The participants were instructed to perform a complete range of motion with each lift, approximately 180° at full extension to maximal flexion. The participants performed two warm-up sets with increasing weight and three minutes of rest between each set. Once the warm-up sets were complete, the participant completed only one repetition of a particular weight. If the attempt was completed without any assistance, the weight was increased, incrementally, and another attempt was performed after the participant was able to rest for 1 minute. The test was terminated once the participant was not able to complete a lift with a given weight, and the last completed lift was recorded. Post-training strength measures were collected two days after the last bout of training but included the same protocols used at baseline.

### Muscle Size Measures

Baseline CSA and VOL measures via MRI were examined before and 24-48 h after the MVC or 1RM measures to prevent any acute testing effects. Post-training measures were obtained 48-96 h after the last bout of training. Prior to the MRI, the participant’s maximum arm circumference was measured with the arm abducted and the biceps maximally contracted at 90°. The peak circumference was the point of measure (POM) marked for MRI. To determine the CSA and VOL, fifteen 16-mm contiguous axial slices with 0 mm interslice spacing were taken from the arm. Scans for both arms were taken by Fast Spoiled Gradient Recalled and Fast Spin Echo with TE 1.9/TR 200 msec. For CSA, a single transverse MRI scan through the POM was used and analyzed via a custom MATLAB (The MathWorks, Natick, MA) program. Rapidia (INFINITT Inc, Seoul, Korea), a PC based software that allows the semi-automatic quantification of muscle, bone and subcutaneous fat was used to analyze volume (17). Whole-muscle VOL was calculated from the summation of six transverse scans taken superiorly and inferiorly to the POM (17).

### Unilateral Resistance Training

The participants performed supervised, progressive strength training, starting no later than 14 days after their pre-training assessments. Subjects 12, 8, and 6 repetition loads were estimated from their pre-training 1RMs using standard percentage tables. The exercises were only performed with their non-dominant (ND) arm and consisted of preacher curls, concentration curls, standing curls, overhead triceps extensions, and triceps kickback. Progression involved decreasing the reps but increasing the weight. Repetition schemes looked like the following: 1) weeks 1-4, 3 sets of 12, 2) weeks 5-9, 3 sets of 8, and 3) weeks 10-12, 3 sets of 6. Weights were increased if subjects could complete the sets easily. The training sessions lasted between 45-60 minutes, alternating between biceps and triceps work. More detailed descriptions of the training protocols were previously published (15, 16).

### Statistics

Statistics were performed with IBM SPSS version 28. There were a total of 1233 participants; however, not every participant had a complete data set, requiring their exclusion on relevant statistical analyses. No data were imputed. Means and standard deviations were ascertained for all variables. For variables not impacted by time (i.e., age and height), pairwise exclusion was utilized. Listwise exclusion was used for the other variables. Factorial repeated measures analysis of variance (RMANOVA) was utilized to determine the whole cohort training effect (i.e., main effect of time) along with the influence of sex. For within sex training effects, RMANOVA was utilized. For between group differences on variables not influenced by time, one-way ANOVA was used. Pearson linear correlation coefficients tested relationships between baseline strength measures (MVC and 1RM) and baseline VOL as well as between percent change of the strength measures and percent change of VOL for the whole cohort and for each sex. Fishers r to z transformations of the correlation coefficients were used to determine statistically significant differences amongst the correlation values. Statistical significance was set at p < 0.05.

### Causal Modeling Protocol and Statistics

To further explore relationships among strength and size variables, we employed causal modeling analytics to our data (18). Measurements for causal modeling including demographics, muscle size (VOL) and muscle strength (1RM and MVC) were collected before and after training. We constrained the cohort to the participants with all variables measured at baseline and 12-week, thus our causal analysis dataset included data from 600 participants (229 males; 371 females) and 22 variables. These 22 variables were categorized into three tiers: (1) *constitutional variables*: sex, age, race; (2) *baseline variables*: 1RM in each arm, body weight, height, systolic and diastolic blood pressure, MVC in each arm, VOL in each arm and, (3) *variables represents change from baseline to 12 weeks* (12 weeks value minus baseline value): 1RM change in each arm, MVC change in each arm, VOL change in each arm and change in body weight. We constrained the model by preventing causal relationships going from variables in a higher tier to variables in a lower tier (i.e. variables in a higher tier cannot <u>cause</u> variables in a lower tier), since variables in the lower tier represent information from earlier time period. To model the qualitative causal structure among the variables, we applied the GFCI (Greedy Fast Causal Inference) algorithm (19). The GFCI can be conceptualized as two sequential stages. During the first stage, the GES (greedy equivalence search) algorithm is applied to estimate the causal relationships among the variables assuming no hidden variables. Then, during the second stage, the FCI (fast causal inference) algorithm is applied to identify the presence of hidden common causes. The GFCI algorithm estimates the presence of causal relationships and hidden common causes up to statistical equivalence represented as a PAG (partial ancestral graph) (19). A PAG contains the follow possible edge types: (1) X → Y: X causes Y, and Y does not cause X; (2) X ↔ Y: hidden variable(s) causing both X and X o→ Y: X→ Y and/or X ↔ Y; (4) X o–o Y: X→Y, and/or Y→X, and/or X ↔ Y. (5) X—Y: X→Y, or Y→X (20). To assess the stability of the discovered causal relationships, we conducted 100 bootstrap resampling of the original dataset and applied the GFCI to each of the bootstrap dataset. The percentage of time an edge is discovered in the bootstrap datasets was computed and represents the stability of the discovered causal relationship. To estimate the qualitative causal relationship among variables (i.e., causal effects), we built Structure Equation Models based on the local estimated PAG (19) around the change muscle strength. The causal effects were estimated with regression models, where the change in muscle strength is the dependent variable, and its causes identified by GFCI are the independent variables.

## Results

### Participant Characteristics

Table 1 contains the participant characteristics for the whole cohort, while Supplemental Digital Content Table 1 contains sex- and trait-specific sample sizes. The whole cohort demonstrated small mean increases in body mass (0.4 ± 2.1 kg) from baseline to post-training, with no sex difference in weight gain. Males were significantly taller and heavier than females at baseline and post-training. However, change in body mass was not significantly different between sexes (p = 0.96) nor was there an age difference between groups (p = 0.07).

**Table 1.**
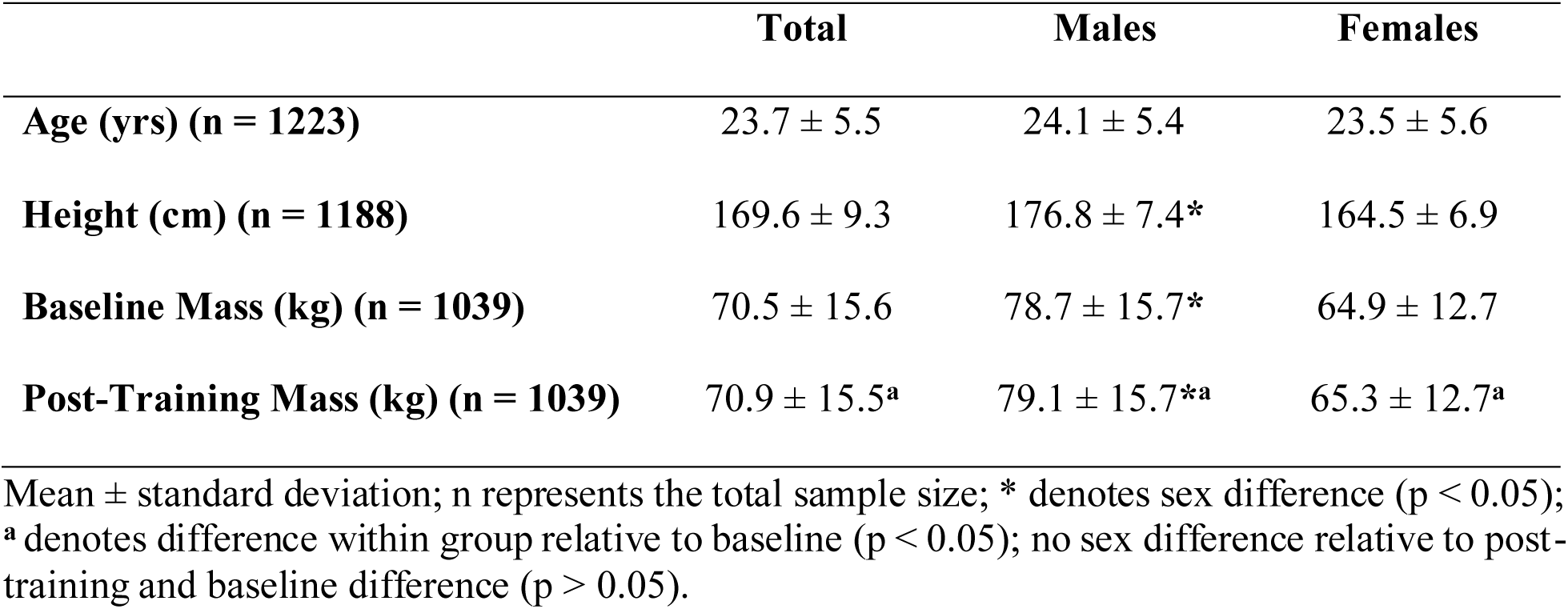
Participant Characteristics.

### Baseline and Post-Training Muscle Strength and Size

Table 2 details baseline and post-training values for strength and size variables. These results include all subjects from our previous report (15) plus a significant number of subjects and volumetric data added. The trained arm saw significant increases in MVC (7.3 ± 7.5 kg), 1RM (3.9 ± 1.9 kg), CSA (5.9 ± 4.4×10^2^ mm^2^) and VOL (5.4 ± 4.1×10^4^ mm^3^) after 12 weeks of unilateral resistance training. When separated by sex, this pattern held true for both males (MVC: 9.3 ± 8.9 kg; 1RM: 4.4 ± 2.1 kg; CSA: 8.1 ± 4.5×10^2^ mm^2^; VOL: 7.5 ± 4.2×10^4^ mm^3^) and females (MVC: 5.9 ± 6.0 kg; 1RM: 3.6 ± 1.7 kg; CSA: 4.6 ± 3.8×10^2^ mm^2^; VOL: 4.3 ± 3.6×10^4^ mm^3^). These strength and size (CSA only) changes are like those we previously reported (15).

**Table 2.**
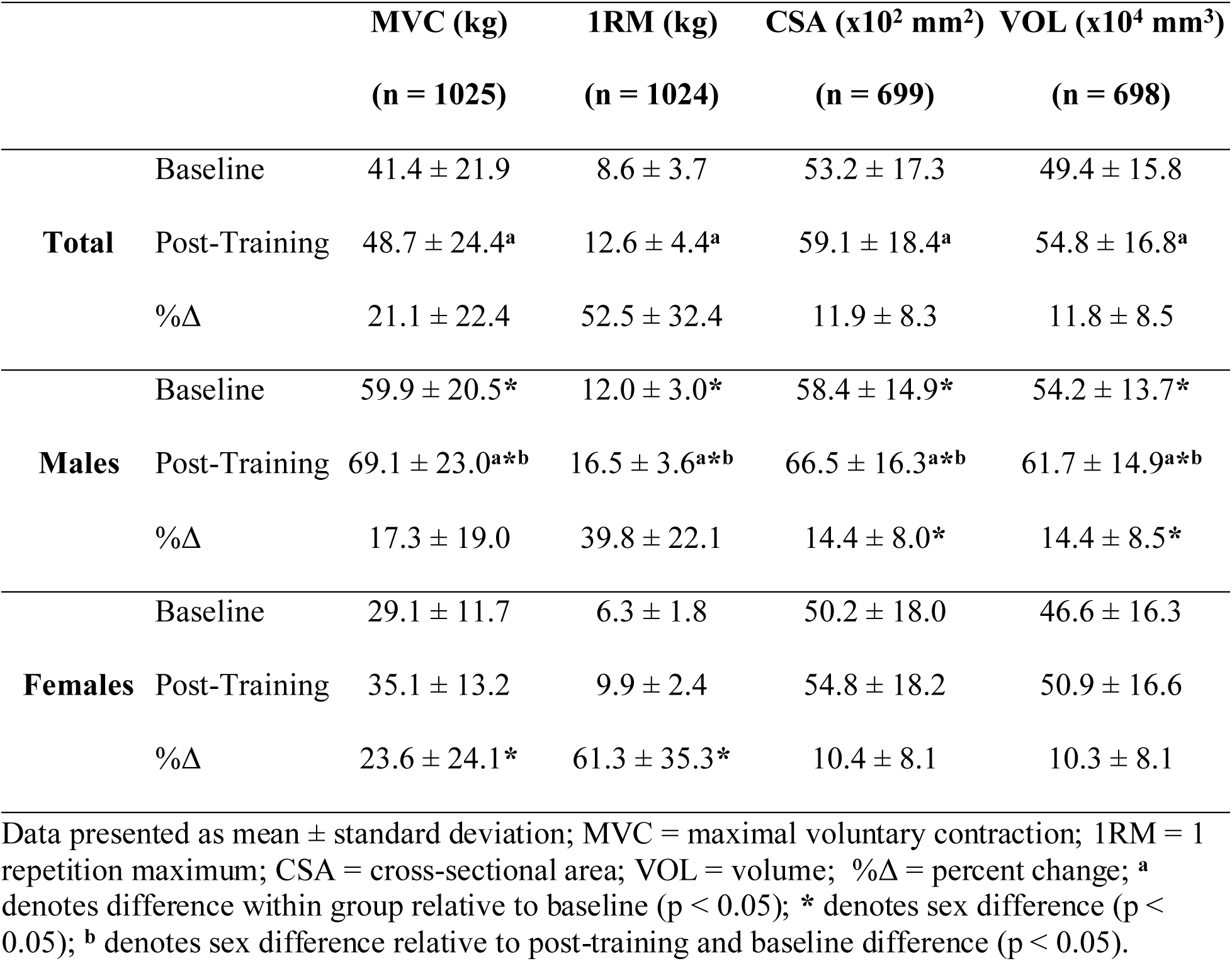
Non-dominant Arm Baseline and Post-Training Values.

Males demonstrated significantly higher mean baseline values for MVC (30.8 kg), 1RM (5.7 kg), CSA (8.2×10^2^ mm^2^), and VOL (7.6×10^4^ mm^3^). These values remained significantly higher post-training (MVC: 34.0 kg; 1RM: 6.6 kg; CSA: 11.7×10^2^ mm^2^; VOL: 10.8×10^4^ mm^3^). While males did demonstrate significantly greater absolute increases in MVC and 1RM, females demonstrated significantly greater relative increases in these variables compared to males (6.3% and 21.5%, respectively) (Table 2). However, males demonstrated significantly greater absolute and relative increases in CSA and VOL (4.0% and 4.1%, respectively). Each of these results also align with our previous report (15).

### Size and Strength Relationships at Baseline and After Training

Baseline non-dominant (Figure 2A) and dominant (Figure 2B) elbow flexors MVC were significantly correlated with baseline non-dominant and dominant whole muscle VOL, despite the relatively small correlation values. The same pattern exists when males and females were tested independently. 1RM findings were similar to MVC patterns. Baseline non-dominant (Figure 3A) and dominant (Figure 3B) elbow flexors 1RM demonstrated small, yet significant correlations with baseline non-dominant and dominant whole muscle VOL. This pattern was also seen independently in males and females.

**Figure 2.**
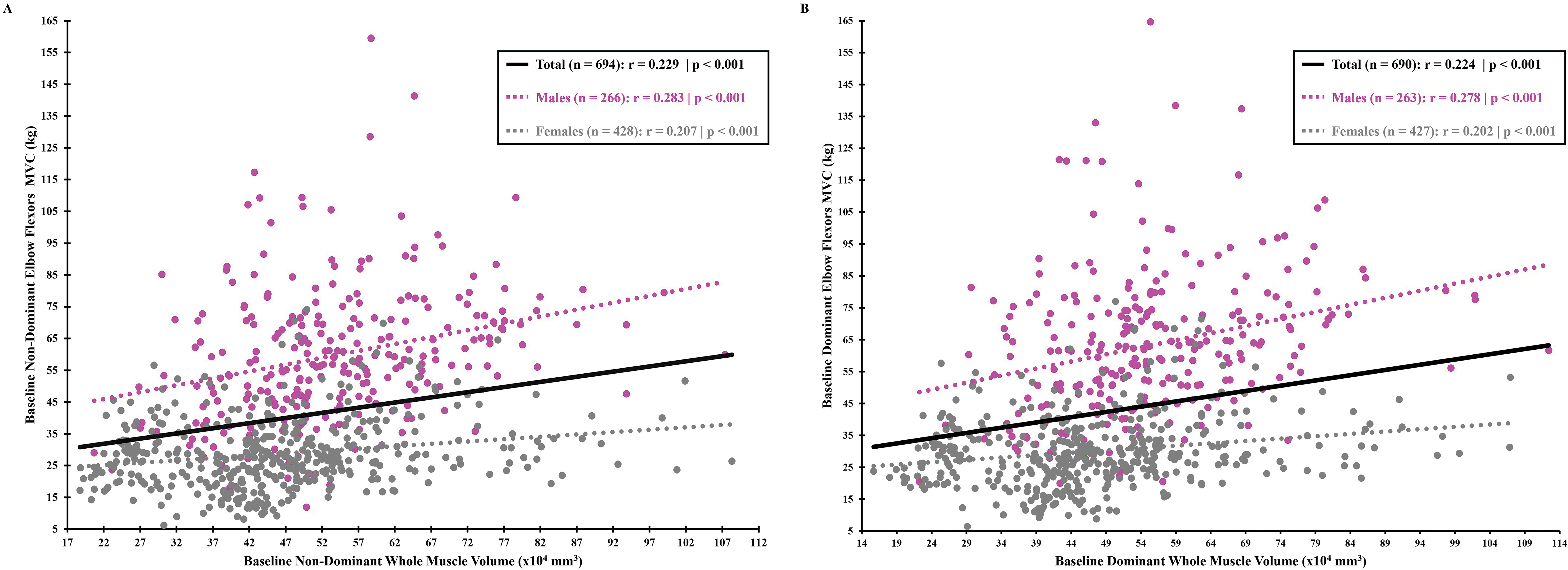
Baseline non-dominant (A) and dominant (B) elbow flexors maximum voluntary contraction (MVC) and whole muscle volume relationship. Solid black line represents the relationship for the total sample when covarying for sex. Dotted gray and magenta lines represent the relationship for males and females, respectively.

**Figure 3.**
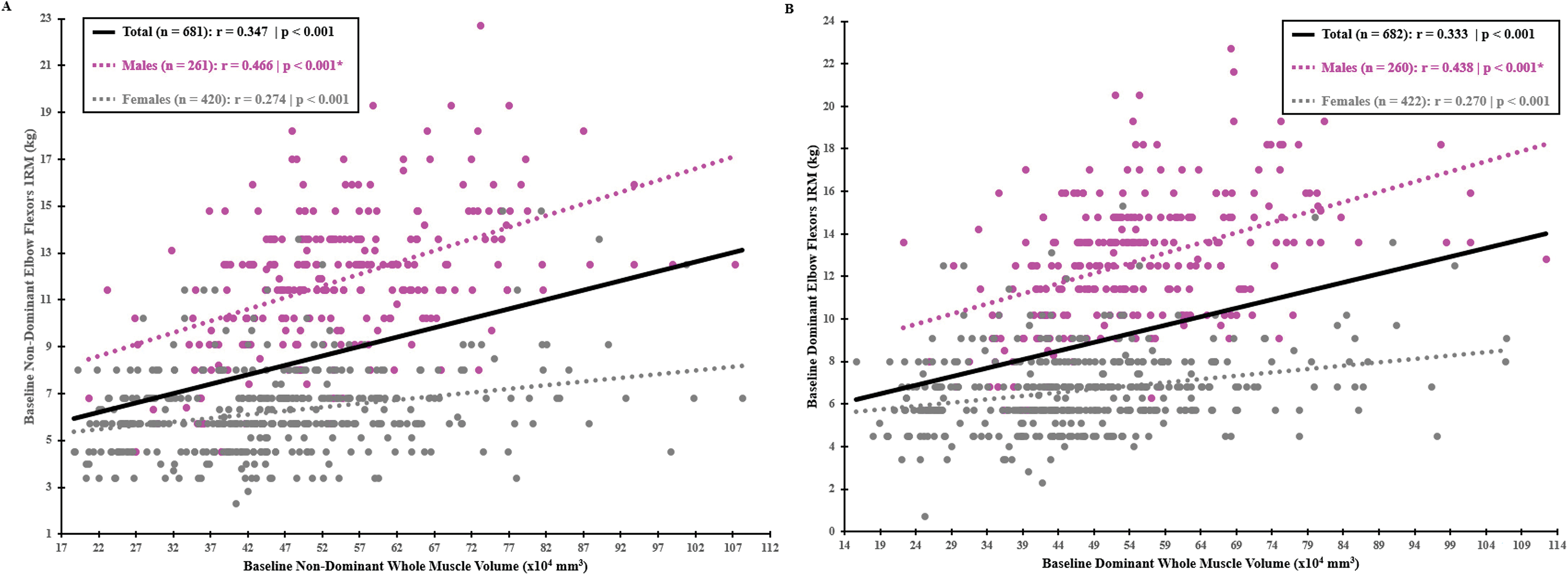
Baseline non-dominant (A) and dominant (B) elbow flexors 1-repetition maximum (1RM) and whole muscle volume relationship. Solid black line represents the relationship for the total sample when covarying for sex. Dotted gray and magenta lines represent the relationship for males and females, respectively. * Denotes significant sex difference for correlation values (p < 0.05).

Size and strength relationships post-training were similar to that seen at baseline (Supplemental Digital Content Figure 2A and 2B). When compared to baseline, there were no differences for the correlation of MVC and VOL for the whole cohort (p = 0.13), males (p = 0.49), or females (p = 0.45). There were also no differences between the correlation values at baseline and post-training for 1RM and VOL for the whole cohort (p = 0.10), males (p = 0.24), or females (p = 0.48).

### Strength and Size Relationships in Adaptation to Training

Correlations for percent changes (% change) in MVC and VOL and 1RM and VOL, respectively, are illustrated in Figures 4A and 4B. MVC % change and VOL % change (Figure 4A) showed a small but significant correlation in the whole cohort (p = 0.001) and in females (p < 0.001), but not in males (p = 0.07). For 1RM % change and VOL % change (Figure 4B), no significant correlations were seen for the whole cohort (p = 0.97), males (p = 0.10), or females (p = 0.11).

**Figure 4.**
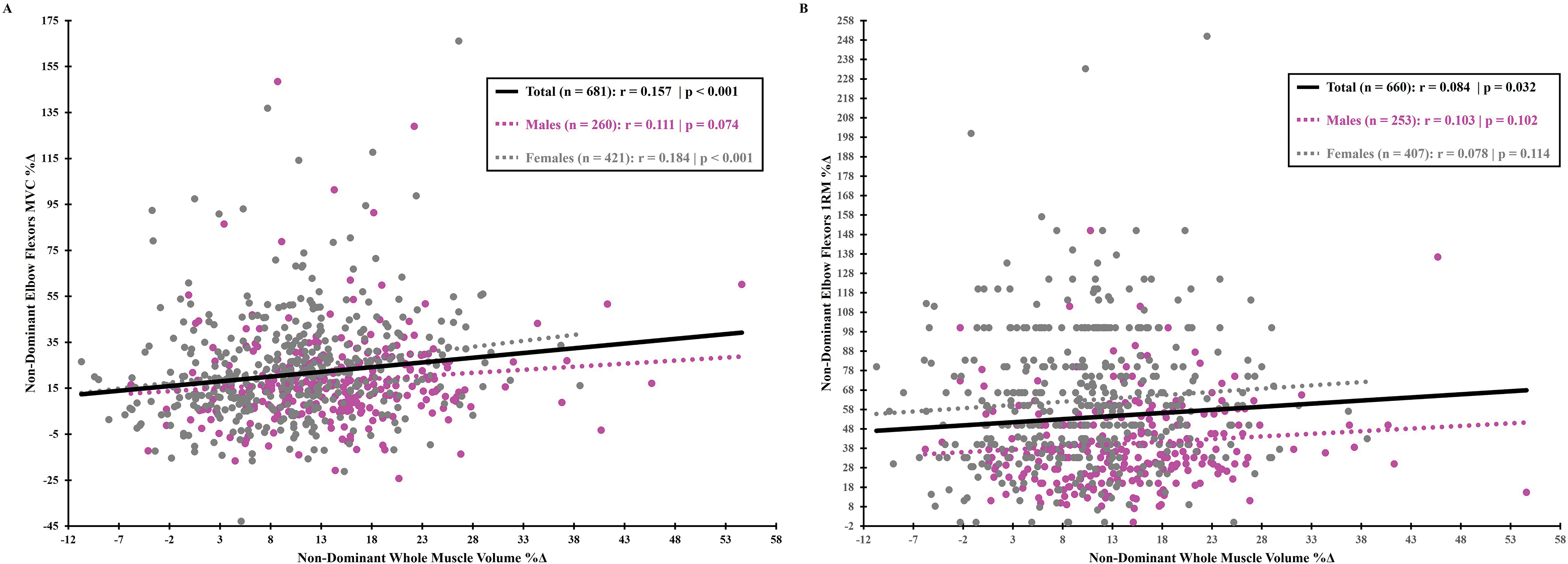
Non-dominant elbow flexors maximum voluntary contraction (MVC) percent change (%Δ) and non-dominant whole muscle volume %Δ relationship (A). Non-dominant elbow flexors 1-repetition maximum (1RM) %Δ and non-dominant whole muscle volume %Δ relationship (B). Solid black line represents the relationship for the total sample when covarying for sex. Dotted gray and magenta lines represent the relationship for males and females, respectively.

### Causal Modeling of Measured Variables

In 22 variables, causal discovery analysis identified 21 edges with 10 representing causal relationships, 3 representing confounded relationship, and 8 representing potentially confounded relationships. Supplemental Digital Content 3 displays the stability of causal relationships that were discovered by the GCFI assessed with bootstrapping. From that table, we were able to derive Figure 5 which demonstrates a simplified estimated causal graph representing the local causal relationships around VOL change and 1RM change. The causal graph indicates that the change of VOL caused the change in 1RM. Also, age is a potential cause of 1RM, but this relationship is potentially confounded. These discovered relationships are fairly stable, appearing 67% and 52% of the times in the 100 bootstrap analysis. Assuming the causal relationship between age and 1RM is not confounded, the estimated causal effect for VOL change on 1RM change was that for every 100,000 mm^3^ increase in VOL, there was a 2.53±0.39 kg increase in 1RM. When accounting for VOL change, the estimated causal effect for age on 1RM change was that for every yearly increase in age, there was a 0.16±0.03 kg decrease in 1RM. The percent variance explained in the 1RM change by the two variables was relatively small, with R^2^ = 0.10.

**Figure 5.**
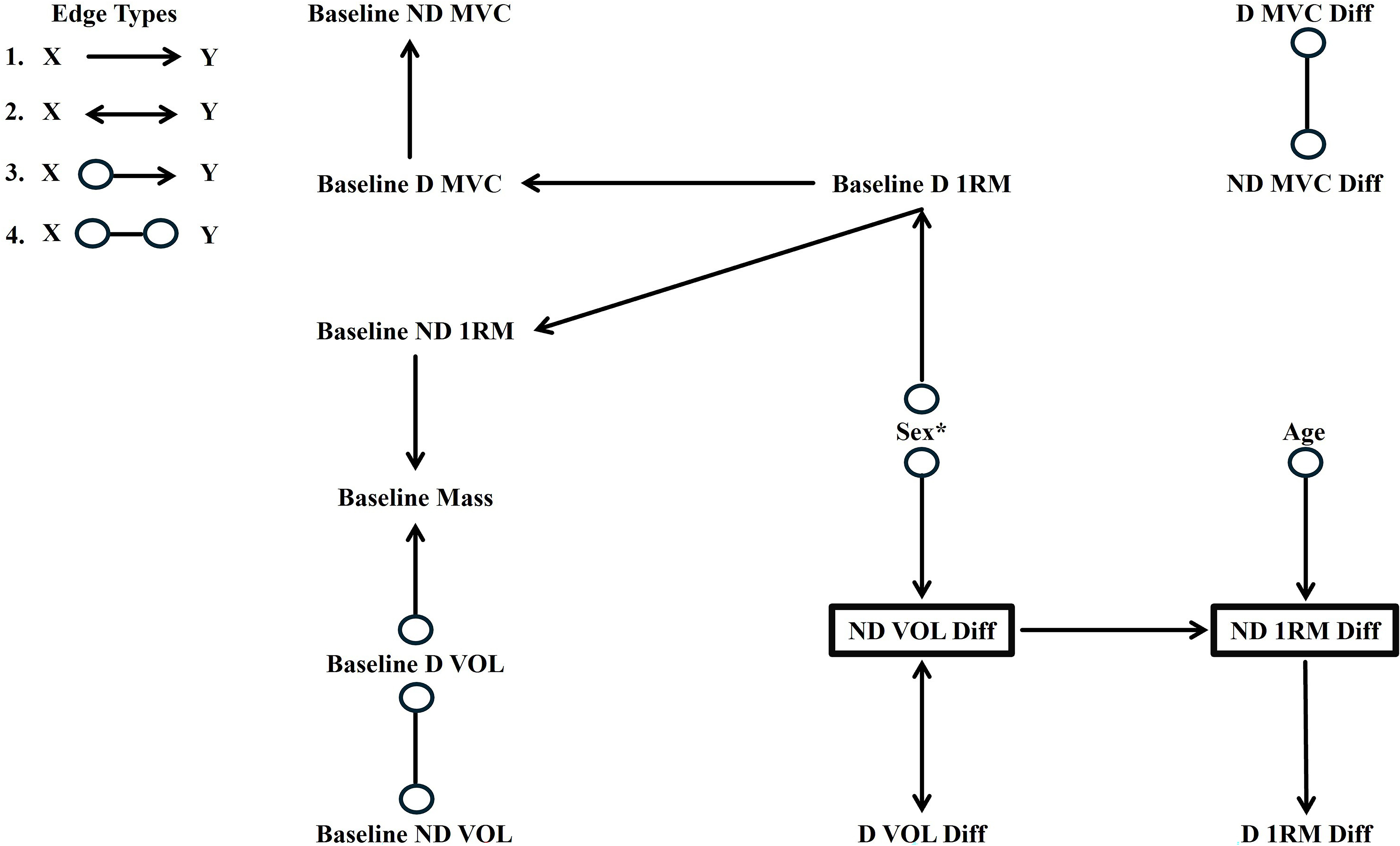
Estimated causal modeling graph. Edge types: Option 1 demonstrates that X is a direct cause of Y or an indirect cause that may include other measured variables or there may be an unmeasured confounder of X and Y. Option 2 demonstrates that there is an unmeasured confounder (L) of X and Y and there may be measured variables along the causal pathway from L to X or from L to Y. Option 3 demonstrates either Option 1 or Option 2. Option 4 demonstrates either of the following holds true: Y is a cause of X, Option 1, Option 2, Y is a cause of X and Option 2, or Option 1 and 2.

## Discussion

This study explored correlation and causal relationships between muscle strength and size before and after 12 weeks of unilateral resistance training in a robust cohort of healthy young adults. We hypothesized that strength and size would demonstrate small, yet significant correlations at baseline and with training, and that sex differences would modify strength/size relationships. The hypothesis regarding strength and size relationships at baseline held true, with significant correlations (despite low correlation coefficients). Interestingly, correlation magnitudes at baseline and post-training were stronger for men than for women (for dynamic but not static strength), hence the overall cross-sectional cohort relationship seems largely driven by male data, inflating a lower relationship magnitude for women. Regarding strength and size changes following resistance training (i.e., adaptations), strength/size relationships become more varied, with only the whole cohort and the females demonstrating significant correlations for static strength and volume, a finding absent in males. The dynamic strength changes, however, were not significantly related to muscle volume changes, highlighting the more complex nature of the 1RM measure.

This study serves as a follow up to our previous exploration of variability and sex differences within the individual traits of muscle size and strength (15), which demonstrated large degrees of inter-subject variation and sex differences in strength and size responses to resistance training in 585 subjects. The previous study used cross-sectional area as its size measure while the current study reports both CSA and 3D muscle volume. Despite the larger sample size in the present study, both analyses find similar CSA sex-dependent changes, with men outpacing women for both absolute and relative gains. Here, we add data supporting greater gains for men in muscle volume. Furthermore, both analyses showed females experience significantly larger increases in both strength measures, despite smaller absolute gains. The current analysis expands our previous analyses to provide insight into strength and size relationships – given that variation exists on both sides, we inquired as to whether big gains in size would predict gains in strength, or vice versa. To that end, we also used a causal modeling approach with our dataset, finding interesting evidence of size and age influences on strength.

### Single Time Point Strength/Size Correlations

Unsurprisingly, baseline strength and volume measures for the whole cohort and subsets were significantly correlated as supported by several previous studies (13, 14, 21, 22). Strength and size were also significantly related after training at similar rates. However, the low correlation coefficients (0.2 to 0.28) highlight that this relationship is weaker than some might think and are different by sex. It is well established that men and women differ in aspects of muscle size and strength, both at baseline and following training (15). Here, we show that the baseline correlation values for 1RM and VOL were significantly larger in males compared to females, elevating the relationship strength in the combined group. Males exhibited significantly higher absolute strength and size values, explained, in part, due to males demonstrating ∼40% greater skeletal muscle mass in the upper body in comparison to females (23). Furthermore, while only comparing small numbers of men and women, Sale et al. (24) and Miller et al. (25) demonstrated men to have significantly larger biceps brachii CSA and mean fiber area within the bicep’s brachii and significantly larger type I and type II fibers. Nonetheless, neither study demonstrated differences in fiber type composition (24, 25).

The current study indicates the strength of relationships between strength and size are dependent on mode of strength assessment – with greater sex differences in dynamic strength measures. These differences could be due to structural differences like greater flexibility at the elbow in females (26), making it possibly more taxing on the elbow flexors than a static test at 90 degrees elbow flexion. Another possibility is overall cultural differences in exposure to the sexes to dynamic types of movement as youth – our subjects were all untrained within the last year but may have experienced some sort of exercise training as youth, with that likelihood being increased in males.

### Strength/Size Correlations with Training

While relationships between strength and size at baseline is informative, relative changes in strength and size are most relevant to the pursuit of gains through resistance training. We saw fewer differences between total cohort versus subgroup relationships, with the total cohort and the males demonstrating lower correlations than were present in absolute baseline or post-training data. As both strength and size changes are known to be multifactorial, this pattern was expected and was influenced by the skill related to the strength task. The isometric task is more constrained than the dynamic task, explaining in part why only MVC change was significantly related to VOL change, whereas no such relationship was seen for 1RM change (Figure 4A and B). The static strength relationship was also seen in females but became insignificant in males. This contradicted the findings of both Erskine et al. (13) (in elbow flexors) and Balshaw et al. (14) (in knee extensors), who demonstrated r values of 0.527 and 0.461, respectively. However, the males’ correlation value for the present study was trending towards significance (p = 0.07). The finding for the females agrees with a study by Higbie et al. (12), but the latter study had separated their females into a concentric training-only group and eccentric training-only group, possibly confounding results. Nevertheless, the present study suggests that hypertrophy is weakly tied to increases in static strength.

Dynamic strength measures present different results than our static strength measure. Because of the lack of relationship regarding 1RM change and VOL change, it seems that hypertrophy has no bearing on increasing dynamic strength, at least with the intervention parameters of the current study. This finding contradicts a study by Ahtiainen et al. (27) who demonstrated a significant, but also small correlation of 0.157 for a whole cohort of 283 individuals. However, this study had mixed methods for assessing changes in muscle CSA, smaller representative sample of females (only ∼36% were female), and an older population (∼48% were 45 years or older); all of which could have impacted the results. Nevertheless, practically speaking, changes in 1RM are more applicable to the training individual than changes in MVC when it comes to traditional resistance training, and it seems that no matter the sex, an individual can get stronger without experiencing hypertrophy.

### Possible Biases from Combining Sexes and Sample Size Differences

We focus here on possible sex differences in our outcomes of interest and analyze outcomes together and separately by sex. Our results may be partially explained by inherent sex differences, but also by potential bias in how we are looking at the data. Most exercise training studies suffer from low statistical power given fairly low sample sizes, which become even smaller in subdividing by sex. Here, we have a very large cohort with the ability to see if sex-independent cohorts follow the patterns of the combined cohort. For example, the MVC and VOL change correlation values (Figure 4A) for the whole cohort (0.16) are larger than the male correlation value (0.11) and more aligned to that of females (0.18). Similarly, the combined group is significantly correlated for 1RM and VOL but neither of the two constituent groups are significantly related.

One possible explanation for these findings is bias from Simpson’s paradox, or more specifically, the Amalgamation Paradox (AMP) (28). This mathematical tenet states that relationships among collated data (such as combining men and women) could be misleading, demonstrating the need for caution when interpreting the data as a whole cohort rather than by subset (in our case, by sex). While the directionality of the correlations here does not reverse, which can happen in the paradox, the magnitude of the relationship is different by sex and this, at times, can flip the results in a test of significance.

Another possible confounding factor in our data is sample size changes when sexes are separated for analyses. Given the weak nature of our strength/size relationships, our tests for significance of the relationship are influenced heavily by our population size. A primary strength of the current work is our robust sample size – detailed in Table 1 and Supplementary Digital Content 1 (broken down by sex). Our total population exceeded 1200 subjects, however not all of these subjects finished training, such that most of the strength changes are in ∼1000 total subjects (∼40% male) and size in ∼700 subjects (just under 40% male). When running tests within sex, our sample sizes were between 260-615. We noticed that the significance results varied with these sample sizes, and we wondered whether the full cohort relationships would still be significant at those population sizes. To test this, we ran randomly generated subcohorts of 400 and 250, using the same % sex in each as compared to the whole cohort. These tests can be seen in Supplemental Digital Content 4, demonstrating the influence of sample size on the significance result. As most training studies are nowhere near even the 250 sample size, it is unlikely that a significant relationship in strength and size gains would be seen.

### Volume Changes Cause 1RM Changes with Training

We also applied causal modeling techniques aiming to discover the mechanistic relationships among measured variables. The causal modeling methods allows for the discovery of causal relationships from observational data up to statistical equivalence (18). These techniques have been applied to biomedical data from various domains for knowledge discovery (19, 29). Our causal model suggests that changes in VOL did partly cause changes in 1RM. Age was also estimated as a potential cause for 1 RM, but this relationship could be confounded. It is worth noting that for our correlation analysis we examined the relative difference in the outcomes, but in the causal modeling we examined the absolute difference between the outcomes. These different ways of representing VOL and 1RM changes from baseline assess their relationships through different lenses. In the correlation analysis, we chose to examine relative changes due to the importance of quantifying the ability of a muscle to adapt from its baseline. In the causal analysis, due to the ability to be able to simultaneous model the relationship among many variables, the baseline VOL and 1RM values are explicitly represented in the model to account for potential influence of the baseline values. Though the correlation between the relative changes of VOL and 1RM is not statistically significant, the correlation between the absolute changes of VOL and 1RM is significant (see digital supplement), which is consistent with our causal modeling results.

An important note to make is that while the present study utilized a more traditional hypertrophy program (i.e., 6-12 reps of relatively low loads (15, 30)), there is evidence supporting the use of high loads (>60%) to induce greater strength and similar hypertrophy gains in individuals (30). While it would be interesting to utilize the present study’s causal modeling approach to further investigate high load vs low load programs, the meta-analysis by Schoenfeld et al. (30) and the present study further enhance the idea of prescribing resistance training on an individual basis with a balance between the individual’s goals and needs.

### Limitations

Both strength and size are complex traits, and we could not account for all possible influences on each trait. Nor did we track them at time points during intervention beyond baseline and post-training. For example, a main factor that seems to explain strength increases are various neurological factors, which were not considered in this study, also change in response to resistance training. McArdle, Katch, and Katch (31) suggest that the strength increases resulting from the first two weeks of resistance training are attributable primarily to neurological factors (∼90%) whereas hypertrophy accounts for only 10%. However, afterwards, hypertrophy becomes more important as both hypertrophy and neurological factors completely flip after eight weeks of training. Alongside this, there were only pre- and post-training measures. Had there been additional time points, this could have allowed for better refinement of time-related changes in the measured variables. Furthermore, having neurological factors and additional time points could have better informed our causal model.

Other factors that could have played a role in muscle adaptations were diet changes and the single site for imaging. It is known that skeletal muscle size is influenced by a balance of protein synthesis and breakdown. To get an increase in size, the muscle needs to be synthesizing more protein than it is breaking down and requires an individual to eat an appropriate number of calories and protein (32). Additionally, a recent review (33) demonstrated that addition of muscle may occur at different parts of a muscle (e.g., distal portion rather than middle). While the evidence discussed did not pertain to elbow flexors, it is likely that this concept could hold true. However, the present study only measured VOL around the baseline maximal circumference; thus, ignoring other areas that could have experienced significant growth.

### Conclusions

Here, we leverage a large cohort of young, healthy adults that performed 12-wk of resistance training to 1) quantify the relationship of muscle strength/size and its response to training and 2) to define sex differences in these relationships. We found significant but weak relationships at baseline and post-training that were larger in men than women, especially for dynamic strength. The strength/size relationships are weaker in the adaptation response, becoming non-significant for dynamic strength gains. These data highlight that resistance training responses for strength (especially dynamic strength) and size gains are largely independent from one another, meaning that practitioners should set their goals and training style (i.e., high vs low loads) specifically to the outcomes they hope to achieve. However, our causal models suggest partial influence of muscle size and age on muscle strength. We also highlight the possible confounding of training study results when males and females are combined for analyses.

## Acknowledgements

This study contains data supported by the National Institutes of Health (National Institute of Neurological Disorders and Stroke and National Institute of Arthritis and Musculoskeletal and Skin Diseases) 1R01NS040606 and 9R01AR055100 - Functional SNP’s Associated with Human Muscle Size & Strength (2001-2010). We would also like to thank Nathan Deiwert for making the 1RM and MVC testing images in Figure 1. We have no conflicts of interest to disclose. The results of the present study do not constitute endorsement by the American College of Sports Medicine. All results herein are presented clearly, honestly, and without fabrication, falsification, or inappropriate data manipulation.

## Notes

### Competing Interest Statement

The authors have declared no competing interest.

