## Supplemental Tables and Figures for "Muscle Strength and Size Relationships with Unilateral Progressive Resistance Training"

**Supplemental Digital Content 1.** Sex-specific sample sizes for physical characteristics, strength, and size variables.

|  | **Males** | **Females** |
| --- | --- | --- |
| **Age** | 500 | 723 |
| **Height** | 487 | 701 |
| **Baseline Mass** | 421 | 618 |
| **Post-Training Mass** | 421 | 618 |
| **MVC** | 410 | 615 |
| **1RM** | 415 | 609 |
| **CSA** | 260 | 439 |
| **VOL** | 260 | 438 |

MVC = maximal voluntary contraction; 1RM = 1-repetition maximum; VOL = volume; CSA = cross-sectional area

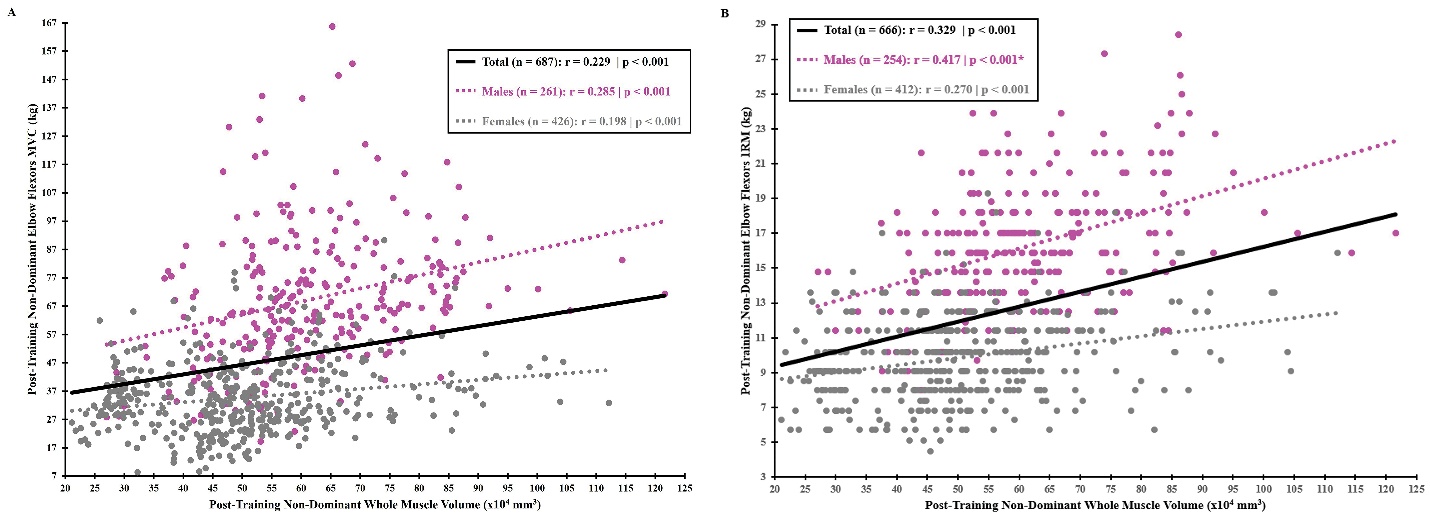

**Supplemental Digital Content 2.** Post-training non-dominant elbow flexors maximum voluntary contraction (MVC) and volume relationship (A). Post-training non-dominant elbow 1-repetition maximum (1RM) and volume relationship (B). Solid black line represents the relationship for the total sample when covarying for sex. Dotted magenta and gray lines represent the relationship for males and females, respectively. * denotes significant sex difference for correlation values (p < 0.05).

**Supplemental Digital Content 3.** Stability of causal relationships discovered by the Greedy Fast Causal Inference (GFCI) algorithm assessed with bootstrapping.

| **Variable X** | **Variable Y** | **→** | **←** | **↔** | **o→** | **←o** | **o-o** | **No Edge** |
| --- | --- | --- | --- | --- | --- | --- | --- | --- |
| Baseline D MVC | Baseline ND MVC | 75% | 5% | 0% | 0% | 5% | 15% | 0% |
| D MVC Diff | ND MVC Diff | 46% | 9% | 1% | 16% | 11% | 17% | 0% |
| Baseline D 1RM | Baseline ND 1RM | 86% | 3% | 4% | 0% | 6% | 1% | 0% |
| Baseline Mass | Height | 48% | 14% | 36% | 1% | 1% | 0% | 0% |
| SBP | DBP | 78% | 4% | 0% | 2% | 0% | 16% | 0% |
| Sex | Baseline D 1RM | 16% | 0% | 9% | 75% | 0% | 0% | 0% |
| Baseline ND VOL | Baseline D VOL | 26% | 16% | 0% | 2% | 0% | 56% | 0% |
| ND 1RM Diff | D 1RM Diff | 84% | 9% | 1% | 0% | 2% | 4% | 0% |
| D VOL Diff | Mass Diff | 18% | 18% | 27% | 0% | 0% | 37% | 0% |
| Sex | Height | 41% | 0% | 18% | 40% | 0% | 0% | 1% |
| D VOL Diff | ND VOL Diff | 26% | 3% | 27% | 42% | 0% | 1% | 1% |
| Age | ND 1RM Diff | 27% | 0% | 0% | 52% | 0% | 0% | 21% |
| Baseline ND 1RM | Baseline Mass | 64% | 8% | 0% | 4% | 0% | 1% | 23% |
| ND VOL Diff | Mass Diff | 39% | 9% | 0% | 0% | 26% | 0% | 26% |
| Sex | ND VOL Diff | 25% | 0% | 0% | 47% | 0% | 0% | 28% |
| Race | Age | 0% | 0% | 0% | 2% | 0% | 66% | 32% |
| ND VOL Diff | ND 1RM Diff | 67% | 0% | 0% | 0% | 0% | 0% | 33% |
| Baseline D 1RM | Baseline D MVC | 49% | 2% | 2% | 0% | 4% | 0% | 43% |
| DBP | Baseline Mass | 33% | 12% | 0% | 9% | 1% | 0% | 45% |
| Baseline D VOL | Baseline Mass | 1% | 15% | 0% | 29% | 0% | 0% | 55% |
| Baseline ND VOL | Baseline Mass | 0% | 26% | 0% | 12% | 0% | 1% | 61% |
| D MVC Diff | D VOL Diff | 12% | 14% | 0% | 0% | 0% | 12% | 62% |
| Race | Sex | 0% | 0% | 0% | 20% | 0% | 16% | 64% |
| Baseline RHR | SBP | 4% | 22% | 0% | 7% | 0% | 3% | 64% |
| Age | Baseline Mass | 4% | 0% | 2% | 25% | 0% | 0% | 69% |
| **Variable X** | **Variable Y** | **→** | **←** | **↔** | **o→** | **←o** | **o-o** | **No Edge** |
| Sex | Baseline D MVC | 8% | 0% | 1% | 17% | 0% | 0% | 74% |
| Baseline D MVC | SBP | 19% | 0% | 0% | 0% | 0% | 5% | 76% |
| Baseline ND 1RM | ND VOL Diff | 23% | 0% | 0% | 1% | 0% | 0% | 76% |
| ND MVC Diff | ND VOL Diff | 7% | 10% | 0% | 6% | 0% | 0% | 77% |
| D 1RM Diff | Mass Diff | 15% | 5% | 0% | 0% | 0% | 3% | 77% |
| Baseline Mass | SBP | 21% | 0% | 0% | 1% | 0% | 0% | 78% |
| Baseline D MVC | D MVC Diff | 11% | 0% | 6% | 2% | 0% | 0% | 81% |
| Baseline D 1RM | SBP | 18% | 0% | 0% | 0% | 0% | 0% | 82% |
| Race | Height | 2% | 0% | 0% | 16% | 0% | 0% | 82% |
| Baseline D MVC | ND MVC Diff | 10% | 0% | 0% | 7% | 0% | 0% | 83% |
| Baseline D VOL | Height | 15% | 0% | 0% | 0% | 0% | 0% | 85% |
| Baseline ND 1RM | Baseline ND MVC | 4% | 5% | 0% | 0% | 4% | 0% | 87% |
| Baseline ND 1RM | D MVC Diff | 10% | 0% | 1% | 1% | 0% | 0% | 88% |
| Baseline D 1RM | D 1RM Diff | 1% | 0% | 11% | 0% | 0% | 0% | 88% |
| Baseline ND MVC | SBP | 10% | 0% | 0% | 0% | 0% | 2% | 88% |
| Sex | SBP | 6% | 0% | 0% | 5% | 0% | 0% | 89% |
| Height | Baseline D MVC | 10% | 0% | 0% | 0% | 0% | 1% | 89% |
| Baseline ND 1RM | Baseline RHR | 10% | 0% | 0% | 0% | 0% | 0% | 90% |
| D 1RM Diff | D MVC Diff | 7% | 3% | 0% | 0% | 0% | 0% | 90% |
| Baseline D VOL | SBP | 6% | 1% | 0% | 3% | 0% | 0% | 90% |
| Baseline ND 1RM | ND MVC Diff | 10% | 0% | 0% | 0% | 0% | 0% | 90% |
| Baseline D MVC | Baseline Mass | 5% | 3% | 0% | 2% | 0% | 0% | 90% |
| ND 1RM Diff | ND MVC Diff | 10% | 0% | 0% | 0% | 0% | 0% | 90% |
| Baseline ND MVC | ND MVC Diff | 1% | 0% | 5% | 4% | 0% | 0% | 90% |
| Baseline D 1RM | Baseline Mass | 5% | 4% | 0% | 0% | 0% | 0% | 91% |
| Baseline ND MVC | D MVC Diff | 0% | 0% | 0% | 9% | 0% | 0% | 91% |
| Sex | Age | 0% | 0% | 0% | 0% | 0% | 8% | 92% |
| **Variable X** | **Variable Y** | **→** | **←** | **↔** | **o→** | **←o** | **o-o** | **No Edge** |
| Sex | ND 1RM Diff | 3% | 0% | 0% | 5% | 0% | 0% | 92% |
| Height | Baseline RHR | 7% | 0% | 0% | 0% | 0% | 0% | 93% |
| Baseline ND 1RM | SBP | 7% | 0% | 0% | 0% | 0% | 0% | 93% |
| Baseline D 1RM | Baseline RHR | 5% | 0% | 0% | 0% | 2% | 0% | 93% |
| Baseline ND MVC | Baseline Mass | 6% | 0% | 0% | 0% | 0% | 0% | 94% |
| ND 1RM Diff | Mass Diff | 5% | 0% | 0% | 0% | 0% | 1% | 94% |
| Dominant Arm | D VOL Diff | 0% | 0% | 0% | 5% | 0% | 0% | 95% |
| Height | ND 1RM Diff | 5% | 0% | 0% | 0% | 0% | 0% | 95% |
| Sex | Baseline ND 1RM | 1% | 0% | 2% | 1% | 0% | 0% | 96% |
| Race | Dominant Arm | 0% | 0% | 0% | 0% | 0% | 4% | 96% |
| Baseline RHR | D MVC Diff | 2% | 0% | 0% | 2% | 0% | 0% | 96% |
| Age | SBP | 2% | 0% | 0% | 2% | 0% | 0% | 96% |
| Age | D 1RM Diff | 0% | 0% | 0% | 3% | 0% | 0% | 97% |
| Race | ND 1RM Diff | 0% | 0% | 0% | 3% | 0% | 0% | 97% |
| Baseline ND VOL | Height | 3% | 0% | 0% | 0% | 0% | 0% | 97% |
| Baseline ND VOL | D MVC Diff | 1% | 0% | 0% | 2% | 0% | 0% | 97% |
| Baseline ND MVC | Sex | 0% | 2% | 1% | 0% | 0% | 0% | 97% |
| Baseline D 1RM | Baseline D VOL | 3% | 0% | 0% | 0% | 0% | 0% | 97% |
| ND 1RM Diff | Baseline D 1RM | 0% | 2% | 1% | 0% | 0% | 0% | 97% |
| Baseline ND 1RM | ND 1RM Diff | 0% | 0% | 2% | 0% | 0% | 0% | 98% |
| Race | Baseline RHR | 0% | 0% | 0% | 2% | 0% | 0% | 98% |
| Race | ND VOL Diff | 1% | 0% | 0% | 1% | 0% | 0% | 98% |
| Baseline ND MVC | Baseline RHR | 1% | 1% | 0% | 0% | 0% | 0% | 98% |
| Age | DBP | 0% | 0% | 0% | 2% | 0% | 0% | 98% |
| Baseline ND 1RM | Baseline D MVC | 2% | 0% | 0% | 0% | 0% | 0% | 98% |
| Baseline RHR | ND VOL Diff | 1% | 0% | 0% | 1% | 0% | 0% | 98% |
| Baseline D 1RM | ND VOL Diff | 2% | 0% | 0% | 0% | 0% | 0% | 98% |
| **Variable X** | **Variable Y** | **→** | **←** | **↔** | **o→** | **←o** | **o-o** | **No Edge** |
| Dominant Arm | Baseline RHR | 0% | 0% | 0% | 1% | 0% | 0% | 99% |
| Baseline ND MVC | D VOL Diff | 1% | 0% | 0% | 0% | 0% | 0% | 99% |
| Age | Baseline D 1RM | 0% | 0% | 0% | 1% | 0% | 0% | 99% |
| Height | Baseline ND 1RM | 1% | 0% | 0% | 0% | 0% | 0% | 99% |
| ND VOL Diff | D MVC Diff | 1% | 0% | 0% | 0% | 0% | 0% | 99% |
| Baseline ND 1RM | Baseline D VOL | 1% | 0% | 0% | 0% | 0% | 0% | 99% |
| SBP | D MVC Diff | 1% | 0% | 0% | 0% | 0% | 0% | 99% |
| Height | Mass Diff | 1% | 0% | 0% | 0% | 0% | 0% | 99% |
| Baseline ND MVC | Height | 1% | 0% | 0% | 0% | 0% | 0% | 99% |
| DBP | Baseline D VOL | 1% | 0% | 0% | 0% | 0% | 0% | 99% |
| Baseline RHR | Mass Diff | 1% | 0% | 0% | 0% | 0% | 0% | 99% |
| Baseline D VOL | ND VOL Diff | 0% | 0% | 1% | 0% | 0% | 0% | 99% |
| Sex | Baseline ND VOL | 0% | 0% | 0% | 1% | 0% | 0% | 99% |
| Baseline ND VOL | Baseline ND 1RM | 1% | 0% | 0% | 0% | 0% | 0% | 99% |
| Race | ND MVC Diff | 0% | 0% | 0% | 1% | 0% | 0% | 99% |
| Baseline ND VOL | ND 1RM Diff | 0% | 0% | 0% | 1% | 0% | 0% | 99% |

Values in cell indicate the percentage of times a specific edge type was discovered between a pair of variables in 100 bootstrap resampling. Percentages that are closer to 100% or closer to 0% percent indicate consistent presence or absence of an edge type and are more stable. The possible edge types: (1) X → Y: X causes Y, and Y does not cause X; (2) X ← Y, Y causes X, and X does not cause Y; (3) X ↔ Y: hidden variable(s) causing both X and Y; (4) X o→ Y: X → Y and/or X ↔ Y; (5) X ←o Y: X ← Y and/or X ↔ Y; (6) X o–o Y: X → Y, and/or X ← Y, and/or X ↔ Y; (7) No Edge. D = dominant arm; ND = non-dominant arm; MVC = maximal voluntary contraction; Diff = post – pre; 1RM = 1-repetition maximum; SBP = systolic blood pressure; DBP = diastolic blood pressure; VOL = volume; RHR = resting heart rate.

**Supplemental Digital Content 4**. Whole cohort and randomized subcohort non-dominant arm strength and size correlations

| **Correlation** | **Whole Cohort** | **400 Subcohort** | **250 Subcohort** |
| --- | --- | --- | --- |
| Pre MVC & VOL | r = 0.229 \| p = <0.001 | r = 0.240 \| p = <0.001 | r = 0.265 \| p = <0.001 |
| Pre 1RM & VOL | r = 0.347 \| p = <0.001 | r = 0.262 \| p = <0.001 | r = 0.309 \| p = <0.001 |
| Post MVC & VOL | r = 0.229 \| p = <0.001 | r = 0.215 \| p = <0.001 | r = 0.221 \| p = <0.001 |
| Post 1RM & VOL | r = 0.329 \| p = <0.001 | r = 0.261 \| p = <0.001 | r = 0.332 \| p = <0.001 |
| MVC %Δ & VOL %Δ | r = 0.157 \| p = <0.001 | r = 0.102 \| p = 0.043 | r = 0.092 \| p = 0.146***** |
| 1RM %Δ & VOL %Δ | r = 0.084 \| p = 0.032 | r = 0.044 \| p = 0.377***** | r = 0.051 \| p = 0.425***** |

MVC = maximal voluntary contraction; VOL = volume; r = partial correlations covaried for sex; 1RM = 1-repetition maximum; %Δ = percent change; ***** denotes **nonsignificant** p-value (p > 0.05).

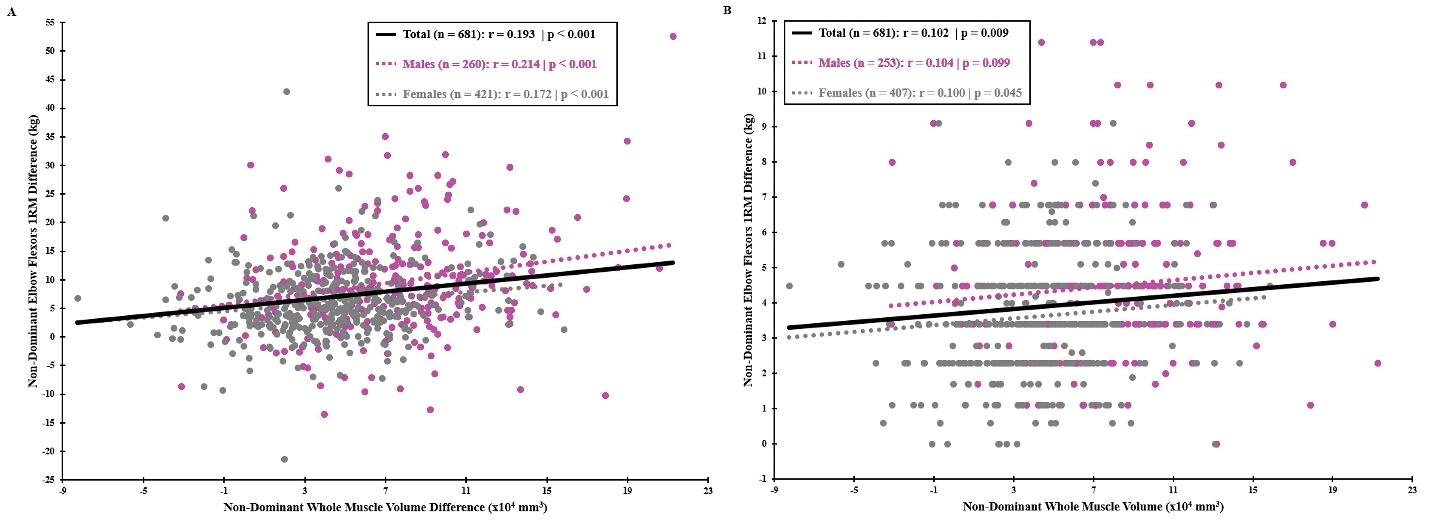

**Supplemental Digital Content 5.** Non-dominant elbow flexors maximum voluntary contraction (MVC) difference and non-dominant whole muscle volume difference relationship (A). Non-dominant elbow flexors 1-repetition maximum (1RM) difference and non-dominant whole muscle volume difference relationship (B). Differences for all variables were calculated as post minus pre. Solid black line represents the relationship for the total sample when covarying for sex. Dotted magenta and gray lines represent the relationship for males and females, respectively.
